# Prevalence and Risk Factors for Child Labour and Violence against Child in Egypt using Bayesian Geospatial Modelling with Multiple Imputation

**DOI:** 10.1101/546697

**Authors:** Khaled Khatab, Maruf A. Raheem, Benn Sartorius, Mubarak Ismail

## Abstract

**Background:** The incidence of child labour, especially across the developing nations is of global concern. The use of children in employment in developing economies constitutes a major threat to the societies, and concerted efforts are made by the relevant stakeholders towards addressing some of the factors/issues responsible for the menace. Significant risk factors include socio-demographic and economic factors, including poverty, neglect, lack of adequate care, and exposure of children to various grades of violence, parental education status, gender, place of residence, household size, residence’s type or size, wealth index, parental survivorship and household size.

**Objectives:** This study, therefore, focuses on identifying socio-demographic/economic and geospatial factors associated with child labour participation.

**Methods:** We utilised 2014 Demographic and Health Survey (EDHS) from the Ministry health and Population in Egypt, with the record of **20, 560** never married children aged 5-17years engaging in economic activities, in and out of their home. The data focused on demographic and socio-economic characteristics of household members. Multivariate Bayesian geo-additive models were employed to examine the demographical and socio-economic factors for children working less than 16hrs; between 16 and 45hrs and above 45hrours weekly.

**Results:** The results showed that at least 31.6% of the children in the age group from 5-10 were working, 68.5% of children aged 11-17 years engaging in child labour (wage), while 44.7 of the children in the age group from 5-10 were engaged in hazard work. From the multivariate Bayesian geo-additive models, the female children (with male children as reference category) working at least 16hrs (OR: 1.3; with 95% CI: 1.2-1.5) are more likely to be engaged in child labour than those working 16 to 45hours (OR: 1; 95% CI: 0.3-1.5). Children born to women without formal education, under non-hazardous jobs, irrespective of the hours spent at work, were mostly exposed to child labour with following percentages: 52.9%, 56.8%, 62.4%, compared to children of mothers with some levels of education. Finally, children that have experienced psychological aggression and physical punishment are mostly exposed to child labour than those without such experience across the job types and hours spent.

**Conclusion:** This study revealed a significant influence of socio-demographic and economic factors on the children labour and violence against children in Egypt. Poverty neglects, lack of adequate care and exposures of children to various grades of violence are major drivers of child labour across the country. North-eastern region of Egypt has a higher likelihood of child labour than most other regions, while children who live in Delta are more engaged in hazard work.

## Introduction

The General Assembly of the United Nations issued in 1989 (Resolution 44/25) a convention for the rights of the children that defines a child as every human being below eighteen years of age. This convention emphasized the need to seek to protect the child from performing any work that could be hazardous, interfere with education, or damage his or her health or physical, mental, spiritual, moral or social development [1]. It also necessitated the Member States to take legislative, administrative, social and educational measures to ensure such protection. In addition, these states are obligated, according to this convention, to set a minimum age for employment, determine an adequate system of working hours and working conditions and impose appropriate penalties to ensure the effective application of these conditions.

The International Labour Organisation (2004) reported in 2004 that in 1999–2000 overall poverty in Egypt stood at 20.2%; with the poverty incidence appears to be highest in urban Upper Egypt (36.33%), followed by rural Upper Egypt (34.68%), but it is lowest in the Metropolitan region (9.01%). At least 12 million people could not fulfil their basic food and non-food needs [2].

Recent statistics indicate that there are around 11 million children (under 15) in Egypt and of these 11 million, around, 1.6 million work as child labour (ages 6 to 15) [3]. In 2014, the World Food Program and the European Union reported that the number of employed minors in Egypt jumped to at least 2.7 million. Most of them work six days a week, an average of 12 hours a day or more. They constitute an additional source of income for their families, who depend on the income to provide for one-third of their expenditures. Approximately 78% are employed in the countryside, and most of them are females. Between 1 and 1.5 million are employed in agricultural labour. Of all working children, 84% live in rural areas while16% in urban areas. Some recent statistics indicated that children who were engaged in child labour have experienced punishment and abuse, largely physical or verbal.[3–4]

Egypt has agreed to the ratification of the present Treaty in addition to many international treaties, which eliminate the criminalized economic exploitation of children, including the international labour treaty No. 138 of 1973, which aimed (long-term) for the total elimination of child labour. Also, Egypt has agreed with the international labour treaty No. 182 of 1999, which came complimentary to the Treaty No. 138, which urges the elimination of the worst forms of child labour initially, with the goal of eliminating child labour. All forms of child labours are included by this agreement which also emphasizes the importance of free basic child education [3]. Despite this, the distribution of the population by age indicates that a relatively high percentage of the population is young: those below the age of 15 years represent about 37.5 % of the total population. Out of this 37.5 %, 7 % of Egyptian children were engaged in child labour in 2009 [2]. Egypt is the fourth highest country in child labour according to the World Bank, 2015 [4]

The major contributing factors to child labour in Egypt are poverty, declining economic conditions, and rising inflation which means that the situation is likely to worsen [3]. The poverty index measures severe health deprivation by the proportion of people who are not expected to survive to age 40. Based on this metric, the 2004 Human Development Report (HDR) submitted that 3.1% (2.2 million people) of the total population of Egypt lives on less than USD 1 per day [3]. Figure 1 shows the risk factors influencing household concerning child labour as reported [5]

**Fig 1.**
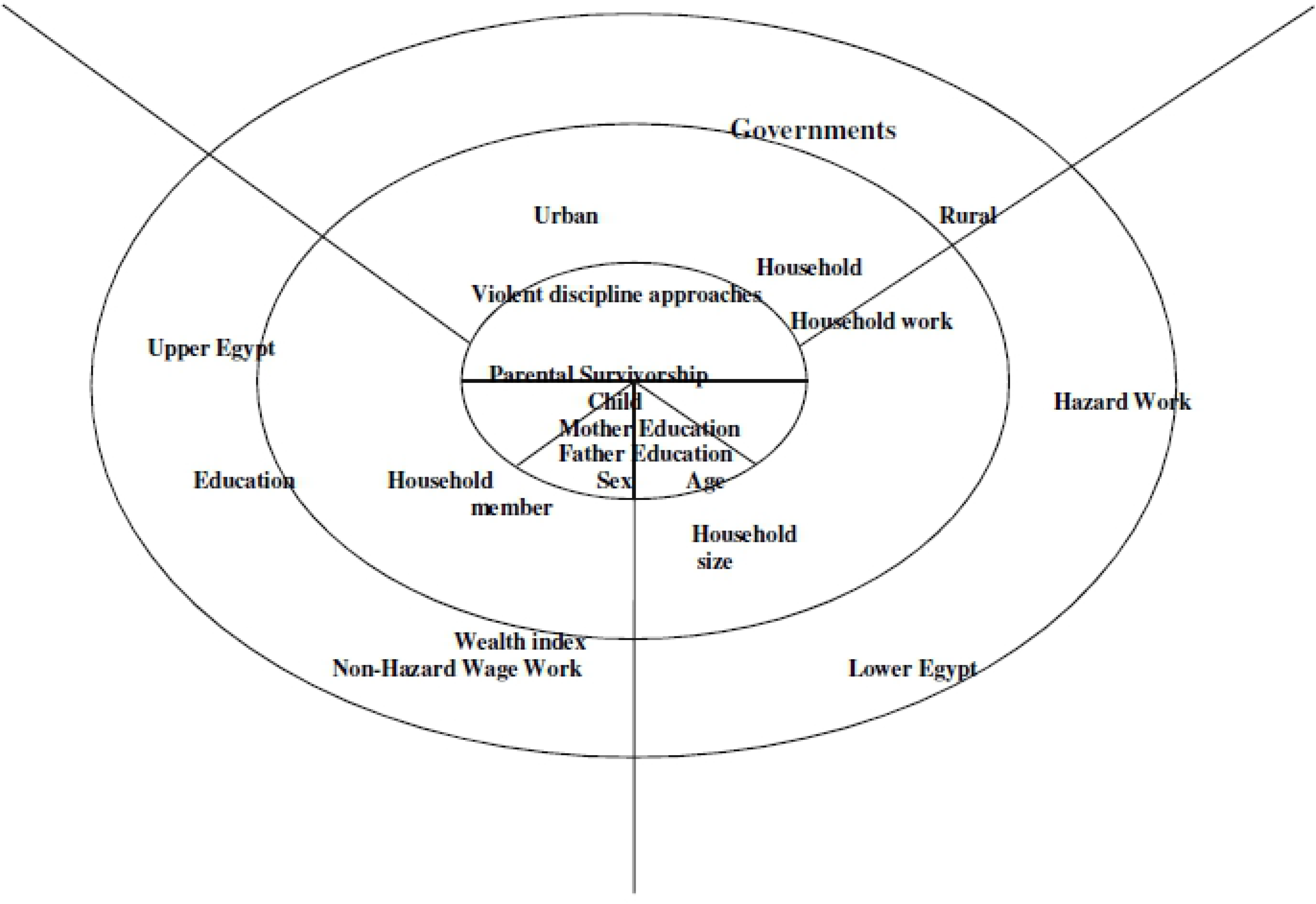
Factors influencing household concerning child labour (Source: Understanding Child’s work, A joint LIO-UNICEF-The World Bank report, 2017).

The Central Agency for Public Mobilization and Statistics (CAPMAS) reported that there are 12 million Egyptians who are homeless, of whom 1.5 million are living in cemeteries [6]. It has been reported also in the same source that the number of people living below the poverty line in Egypt increased to 26.3 % of the population in 2013 compared to 25.2% in 2011 [6]. The report showed that the urban frontier governorates had the lowest poverty rate with 11.4 %, while rural Upper Egypt governorates showed the highest poverty rate with 49.4 %. Also, it was found that among the illiterate, 37 % are poor while only 8 % of those who finished universities were poor. The poor clearly exists in large households with more than 10 members where 67 % of these households are poor [6].

The problem is that it is difficult to find accurate statistics on child workers or accurate studies that have discussed this problem properly. Therefore, there is an urgent need for a study that can describe this problem accurately and discuss the causes of it and provide some suggestions that may help decision makers to handle this issue. In particular, the recent political changes in Egypt may offer a chance to encourage the new government to deal with this problem. Due to the negative effect of this phenomenon on the society, it is important to precisely investigate this problem further and thoroughly using the results to shape the policy and dedicated to improved outcomes for children [7].

## Materials and Methods

The analysis in this work is based on the data obtained from the 2014 Egypt Demographic and Health Survey (EDHS) (**<u>S1 Data</u>**) which is the most recent data on child labour in Egypt. The 2014 EDHS data conducted by Egypt’s Ministry of Health and Population, the National Population Council in collaboration with Macro International. (See Ministry of Health and Population, El-Zanaty and Associates, and ICF International for detail information about methods used in EDHS [8] [9]). As this was a secondary data analysis of open access data, ethical approval was not required. The sample for the 2014 EDHS was designed to provide estimates of population and health indicators, welfare indicators, and other indicators rates for the country as a whole and for six major subdivisions (Urban Governorates, urban Lower Egypt, rural Lower Egypt, urban Upper Egypt, rural Upper Egypt, and the Frontier Governorates).[8]

The sample likewise allows for estimates of most key indicators at the governorate level.

In order to allow for separate estimates for the major geographic subdivisions and the governorates, the number of households selected from each of the major sectors and each governorate was disproportionate to the size of the population in the units.[8]

Thus, the dataset was weighted before proceeding with the analysis.

### Study area and data

The 2014 EDHS on child labour allows for an assessment of several key aspects of the welfare of Egypt’s children. Questions were included on birth registration and living arrangements and the survival status of parents. Data also were collected on the prevalence of injuries and accidents and disabilities among young children. A child’s access to education is critical, and the EDHS obtained information on both the level of pre-school education among young children and children’s participation in primary and secondary school.

The survey also looked at the extent of child labour and at the practices used in disciplining children. In the 2014 EDHS, the first step in the administration of the Child Labour and Child discipline modules involved the identification of a single child age 1-17 years for whom the questions in the modules would be asked depending on the child’s age. If the household included more than one child in the age range, the child for whom the modules were administered was selected using a Kish grid^1^. If the selected child was 1-14 years, the Child Discipline module was administered for the child. To account for the selection of one child per household, the child discipline data are weighted. The weight is based on the de jure population of children age 1-17 years [8].

### Description of outcome variables

The 2014 EDHS considered the never-married children aged 5-17 years who are involved in economic activities inside or outside the home according to the child’s age and a number of hours worked. However, The MICS program has defined thresholds based on the child’s age and the number of hours a child worked during the week to classify children’s involvement in economic activities (MICS2014). This study, however, has used the ILO classifications which are classified based on the number of hours worked per week into three groups as follows: A) Less= 16rs a week; B) between 16 and less than 45hrs a week; and C) over 45hrs a week.

The economic activities were classified into three groups, such as:

A) Non-Hazardous wage work. B) Hazardous wage work. C) Household work.

### Risk factors and covariates

We considered the following socio-demographic factors and the associated risk factors of the child labour as explanatory variables: child’s age (5-17years), sex, household size, place of residence, wealth index, mother education, father education, parental survivorship and violent discipline approaches against children. The wealth index was used as proxies for the socio-economic position of the household because EDHS does not collect information on household income and expenditure. Egypt comprises 27 governorates, which were categorised by EDHS into 7 areas namely: Urban governorates, Lower Egypt urban, Lower Egypt rural, Upper Egypt urban, Upper Egypt rural and Frontier governorates. However, in spatial analysis, we have used 27 governorates to investigate the spatial effects in the prevalence of overlap economic activities of children at the state level. This was achieved using a geo-additive semi-parametric multinomial model [9]. Figure 2 shows the multilevel risk factors of child labour that applied to this study.

**Fig 2.**
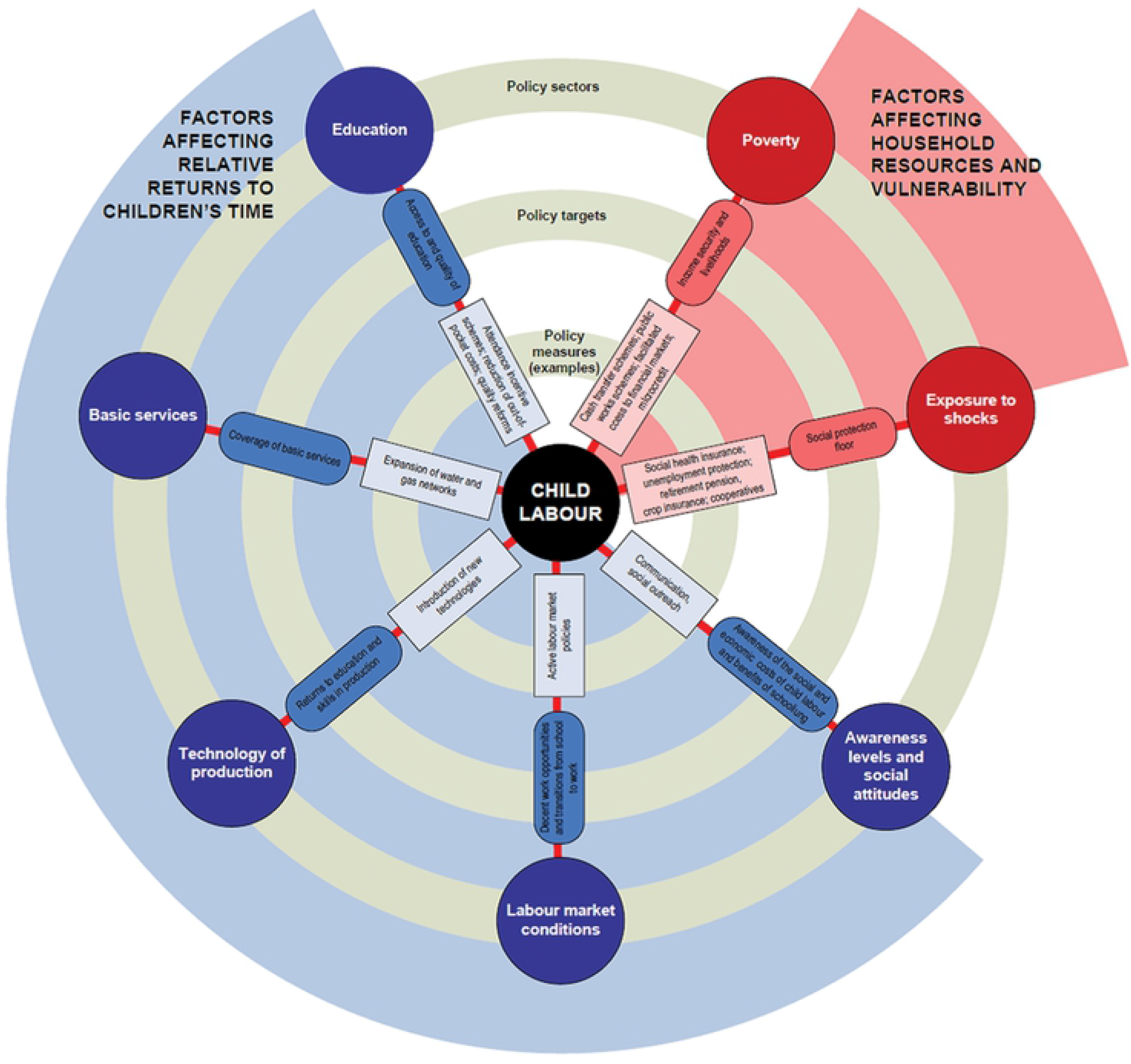
A multilevel risk factors of child labour that applied in this study.

### Statistical Methods

Let *Y_ijk_* and *π_ijk_* represent the type of work and probability of working hours respectively.

Let working hours of at least 15hr diseases be (*k* = 1); working hours falling between 16 and 45hrs a week be (*k* = 2), and the working hours of over 45hrs a week be (*k* = 3).

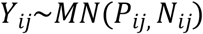

We assume that *Y_ijk_* is distributed as a multinomial distribution, such that: *Y_ij_~MN*(1, *π_ijk_*) where *π_ijk_* = (*π*_*ij*1_,*π*_*ij*2_,*π*_*ij*3_)′. Given some categorical covariates,*Z_ij_*, metrical covariates, *υ_ik_* and state-specific random effect, *S_ik_*, the probability can be modelled thus:

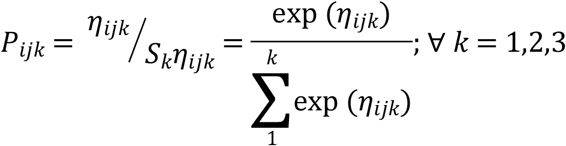

The predictor, *η_ijk_* is given by *η_ijk_* = *z_ij_β_k_* + *f_k_*(*υ_ij_*) + *S_ik_* where *η_ijk_* is a known response function with a logit link function, *β_k_* is the vector of the regression parameters (explanatory variables such as gender, place of residence, etc.) and *f_k_* is a smooth function for the metrical covariates (child’s age) which were assumed to be nonlinear in some previous studies for each of the status categories k [10] [11] [12] [13]. We have included these variables as nonlinear metrical covariates in the early stage of this study; however, the pattern did not show exactly the significance level of each category. Therefore, we used these covariates as linear effects instead to assess the significance level of each category. We set the first category as reference and used the logit link for the modelling.

The random effects, *S_ik_* are district or sub-district specific factors, which can be split into spatially structured variation (*θ_ik_*) and unstructured multinomial heterogeneity (*ϕ_ik_*), such that, *S_ik_* = *θ_ik_* + *ϕ_ik_*. P-spline priors were assigned to the functions; *f*_1_, *f*_2_,…*f_p_* whereas, a Markov random field prior was used for *f*(*S_i_*)[14] [15].

To estimate model parameters, we applied the fully integrated Bayesian approach. Though the estimation method with this model is difficult, the estimated posterior odds ratios (OR) that were produced could be understood as similar to those of normal logistic models. The analysis was carried out using version 2.1 of the BayesX software package, which certifies Bayesian inference, based on Markov chain Monte Carlo (MCMC) simulation techniques [16] [17] [18].

Both the descriptive statistics and chi-square tests were carried out to examine the level of associations between predictors, confounders, and outcome variables using version 14 of STATA; with the p-values of less than 0.05 considered statistically significant. The multinomial logistic regression model was used to determine the degree of associations between the outcome variables (a type of work indicators) and all the predictors. Posterior Odds Ratios (OR) and their 95% confidence intervals (CI’s) were estimated.

### Multiple imputations Analyses (Missing data)

This dataset has a significant proportion of missing data (**S5 Table**) although the missingness is concentrated only in a few variables. At this stage, we will assess the monotone patterns and the joint probability of missing across variables; thereafter which we will identify the potential predictors of each variable that requires modelling ([19], [20], [21], [22]).

One analytic option is to use only that dataset with complete observations, or we can replace the missing values, through a process termed ‘imputation’. The simplest imputation replaces the missing value with mean or median value for that variable; though, this is not a desirable process, especially when one is examining the relationships between variables.

However, the more sophisticated method, termed ‘multiple imputations’, predicts missing values for a variable using existing values from other variables. The predicted values, called “imputes”, are substituted for the missing values, resulting in a full data set called an “imputed data set” [23], [19]. This method imputes dataset using standard procedures for complete data and combining the results from these analyses.

No matter which complete-data analysis is used, the process of combining results from different data sets is essentially the same. In other words, we use the data from units where both (*Y, X*) are observed to learn about the relationship between Y and X. Then, use this relationship to complete the data set by drawing the missing observations from, *X|Y*. This process is completed at if least N=5 times, giving rise to N complete data sets. Then each of these imputed data sets is analysed and we combine the results using specific rules. It does not attempt to estimate each missing value through simulated values but rather to represent a random sample of the missing values [20], [23]. Multiple imputations inference involves three distinct phases as follows:

1. Create imputed data sets which are plausible representations of the data.
2. Perform the chosen statistical analysis on each of these imputed data sets by using standard procedures.
3. The results from the complete data sets are combined “average” for the inference to produce one set of results.

Analyses based on Multiple imputed data will avoid bias only if enough variables that predict the missing values are included in the imputation models. Therefore, including as many predictors as possible tends to make the missing-at-random assumption more plausible [20]. However, including more than 25 predictors will increase the variance explained in the prediction equations [24]

Multiple imputations are a more appropriate choice as a solution to missing data problems as it represents a good balance between quality of results and ease of use. It has been shown to perform favourably compare to other methods in a variety of missing data situations [25], [26]. It can also produce unbiased parameter estimates which reflect the uncertainty associated with estimating missing data. Furthermore, it has been shown to be robust to departures from normality assumptions and provides adequate results in the presence of low sample size or high rate of missing data.

The results of some previous studies using the most commonly used multiple imputation methods, are Expectation Maximization (EM-algorithm) [14], [15] and the Monte Carlo Markov chain (MCMC) [16] [27] method showed there was no significant difference between EM algorithm and MCMC method for item imputation; and that the number of items used for imputation has little impact, either [17].

However, in this study, we have applied MCMC method based on pseudo-random draws and this will allow us to obtain several imputed data. It is known that MCMC can be used with both arbitrary and monotone patterns of missing data. It is known as a collection of techniques for simulating random draws from difficult probability distributions via Markov chains. MCMC also is especially useful in Bayesian statistical analyses and for solving difficult missing-data problems [14].

### Assumptions about Missing Data

If EDHS data contain observations that were missing completely at random, the observations would constitute a random sample of the complete dataset. Multiple imputations assume that the observed variables are predictive of the missing values and that the data are missing at random. Missing data are said to be missing at random (MAR) if the probability that data are missing does not depend on unobserved data but may depend on observed data. MCAR can be viewed as a particular case of MAR.

On the other hand, if the subjects are withdrawn from the study for ethical reasons, missing would not be MAR. This type of missing-data mechanism is called missing not at random (MNAR). For such missing data, the reasons for its missingness must be accounted for in the model to obtain valid results. We looked at missing data patterns and also assessed the extent of missing in the variables that were included in the analysis. Nonlinear relationships were treated using semiparametric models (e.g. generalised additive models (GAMs) [28]. It was important to include the outcome variable (in this case, economic activates) as a predictor in the imputation model because failing to do so will dilute the associations between the outcome and the other variables [18] [19].

## Results

### Prevalence and Sectoral Distribution of Child Labour

**S1 Table** shows statistics on children work and education in Egypt according to the UNESCO Institute for Statistics, 2015. While **S2 Table** shows an overview of children’s work by sector and activity in Egypt according to the US Department of labour report, 2016 [30][31]

**S1 Table** shows that at least. 6.7% of the children in the age group from 5-14 are working, 88.1% of children aged 6-14 years combine schooling with engaging in child labour. Only 10.8% completed the primary school. **S2 Table** provides an overview of children’s work by sector and activity in Egypt. It shows that over 52% of children are working in Agriculture followed by 30.4% who are engaged in service sectors [30][31]

### Distribution of Factors Analysed in Child Labour in Egypt (DHS 2014)

**S3 Table** presents the distribution of socio-demographic factors relating to child labour including violence against children, aged 5-17 years in Egypt with respect to the weekly length of working hours in jobs that are hazardous, non-hazardous and household-based.

The following factors were significantly associated with Non-hazardous wage work (see **S3** Table): gender of a child (P=0.002); residence type, household size, place of residence and wealth indices (P=0.001 each); violent discipline approach, especially: psychological aggression (P=0.002) and physical punishment (P=0.08). However, the following factors: parental education, parental survivor and severe punishment were not significant in this category. For the hazardous wage work, the non-significant factors were household size, Parental Survivorship, and Psychological aggression against children. Gender, the age of children under 2, place of residence, wealth indicators, mothers’ education, physical and severe punishments (each with P=0.001). Father’ education has a slight effect. For household work, all the factors were highly significant (each with P=0.001).

Further, the percentage of male children exposed to child labour is largely higher than those of female children, irrespective of the job type (Non-hazardous, hazardous and household-based work) and the length of jobs, except for female children in a household job, and are working at least 16 hours weekly have the higher percentage of 69.1% and 61.5% respectively than the male children. Also, children from rural communities are more exposed to child labour than their colleagues from urban cities with higher percentages across the board of job type and hours spent at work.

Children living in the medium household and into non-hazardous jobs are apparently having highest exposure rates (53.6%, 50.5% and 50.7%) compared to their mates in both small and large households, irrespective of the hours spent at work. Children from Lower and Upper Egypt were engaged in non-hazardous and household jobs and they have the highest percentage across the three periods of a job than others from another region. However, for the hazardous jobs, children from the Lower Egypt area seem to have the highest percentage of exposure to child labour.

For the wealth indicators, children from the poorest homes were more exposed into non-hazardous jobs with the following percentages (68.4, 53.8 and 44.4) for the three hours of at least 16 hours, 16-45hrs and “above 45hrs” respectively than their colleagues at the remaining wealth indices. The same with the poorest children working at least 45hrs in both hazardous and household jobs have a higher percentage of exposure to child labour than their colleagues from the remaining homes.

Children born to women without formal education, under non-hazardous jobs, irrespective of the hours spent at work, are mostly exposed to child labour with following percentages: 52.7%, 56.8% 62.4%, compared to children of mothers with some levels of education. Also, children whose both parents are alive seem to be most exposed to child labour under household job condition than their mates without living parents.

Finally, children that have experienced psychological aggression and physical punishment are mostly exposed to child labour than those without such experience across the job types and hours spent. The same with those who have experienced severe physical punishment compare to their counterparts who have not experienced severe physical punishment.

### Associate Risk Factors with Child Welfare including Child Labour and Violence against Child issues in Egypt

**S4 Table**, however, displays the multinomial regression results on socio-demographic factors relating to child’s labour, among the children aged 5-17 years in Egypt with respect to the weekly length of working hours in jobs that are hazardous, non-hazardous and household-based. The table presents the estimated effects of the categorical variables: Sex of child, Residence’s type, Household size, Place of Residence, Wealth index, parental education, Parental Survivorship and Violent discipline approaches.

Columns 1&2 present the odds of children working “less than 16 hours” weekly as against those working “between 16 and 45 hours” weekly, under Non-hazardous working condition. For example, the results show that female children (with male children as reference category) working less than 16hrs (OR: 1.3; with 95% CI: 1.2-1.5) are more likely to be impacted by child labour than those working 16 to 45hours (OR: 1; 95% CI: 0.3-1.5). Also, for those under household job, female children working between 16 and 45hours (with OR: 1.7; 95% CI: 0.9-3.2) are more likely to be involved in child labour than those working at less than 16hrs weekly (OR: 1.2; 95% CI: 0.7-2). Finally, under the hazardous working condition, female children working 16-45hrs (OR: 1.6 and 95% CI of 1.4-2.7) are more at risk than those working “less than 16hrs” weekly (OR: 0.6; 95% CI: 0.3-1.5)

Similarly, Children aged 11-15 (OR: 0.5 and 95% CI: 0.4-0.8) are more likely to be at risk than those aged over 15 years (OR: 0.4; 95% CI: 0.3-0.6) under the non-hazardous working condition for “less than 16hrs” and for 16-45hrs of the weekly job. Although, the OR does not seem significant in both age groups. Also, for children under household job, 11-15 of age children are more likely to be at risk of child labour (OR: 2.3 and 95% CI: 0.9-5.6) compared to those who are over 15 and are in 16-45hrs weekly job category. However, for the hazardous job, children aged over 15 years who are working “less than 16hrs” (OR: 1.36; 95% CI: 0.3-3.2) are more at risk of child labour compared to their counterparts who are under age of 15.

Rural children under 16-45hrs of the weekly job are more likely to be at risk than those children working less than 16hrs a week under non-hazardous job. However, urban children under non-hazardous and hazardous job more likely to be at risk compared to rural children, regardless of the number of hours. However, rural children under household jobs are more likely to be affected by child labour compared to urban children.

Children living under medium household who are working 16-45hrs weekly (with OR: 1.4; 95% CI: 1.3-1.6) have increased the risk of child labour in the non-hazardous job than those of hazardous and household jobs. However, for those children in the large household, and working less than16hrs a week under hazardous condition (OR: 1.6; 95% CI: 0.5-2) may likely to have a higher risk, compared to others at the two other working environments.

The poorer children working under the hazardous condition for “less than 16hrs” weekly (OR: 1.93; 95%CI: 0.29-12.8) are likely to be most at risk compared to their colleagues subjected to the two other working conditions. The same experience goes to the middle-class children (OR: 2.3; 95%CI: 0.4-10.07). However, for the children from the richer and the richest families, those under household job and are working “less than 16hrs” weekly, or 16-45hrs are more likely to be at risk of child labour than other children who are in the other two working conditions.

The half-orphans whose fathers are dead tending to be more exposed to the risks of child labour, under non-hazardous and household-based job than those whose mothers are deceased, irrespective of the hours of labour. However, for the children exposed to hazardous jobs, those with deceased mothers are more likely to be at risk than those with deceased father.

Further, children who are exposed to psychological aggression and are working 16-45hrs weekly (OR: 3.7; 95%CI: 1.5-9.5), under non-hazardous job, are more likely to be at risk of child labour than their counterparts subjected to the two other working conditions. The same with children subjected to severe physical punishment. However, the risk likelihood is highest for children in household jobs, who are subjected to physical punishments, especially, those working less than 16hrs weekly (OR: 9.3; 95%CI: 4.2-20.8).

**Figure 3** shows the structured spatial effects of child labour. The results confirmed evidence of regional differences in the likelihood of a child exposed to different type of works. From the graph, it is clear the North-eastern region of Egypt has a higher likelihood of child labour than most other regions. Using Frontier governorate as a reference, children residing in Upper Egypt Rural have the highest risk of having all of three child labours. Hazard work is likely to be in Delta (Suez) Canal, Ismailia, other neighbour cities); whilst, the lower and upper Egypt are more affected by household work. Similarly, the likelihood of being exposed to Non-hazard work is higher in lower and upper Egypt.

**Fig 3.**
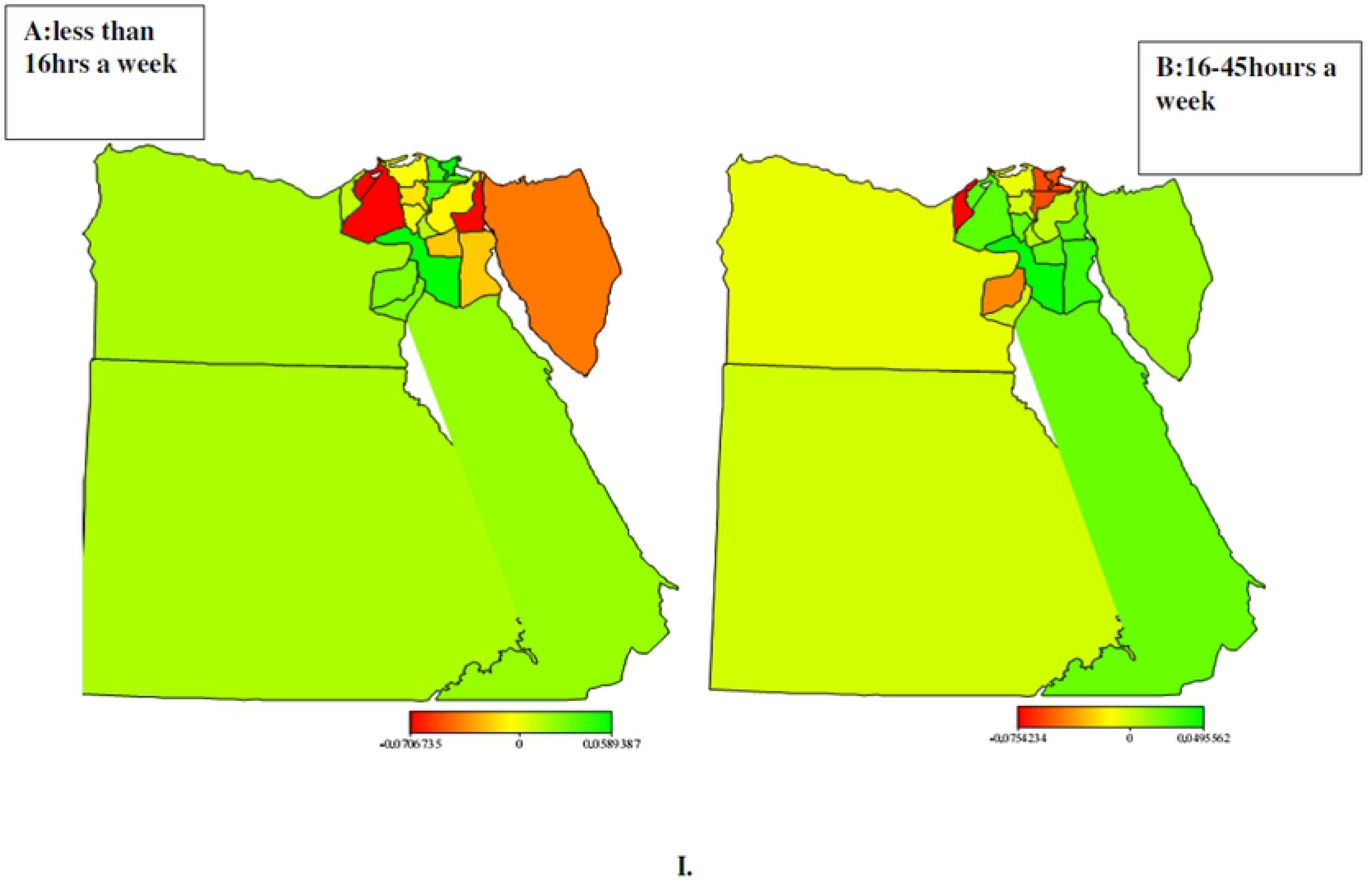
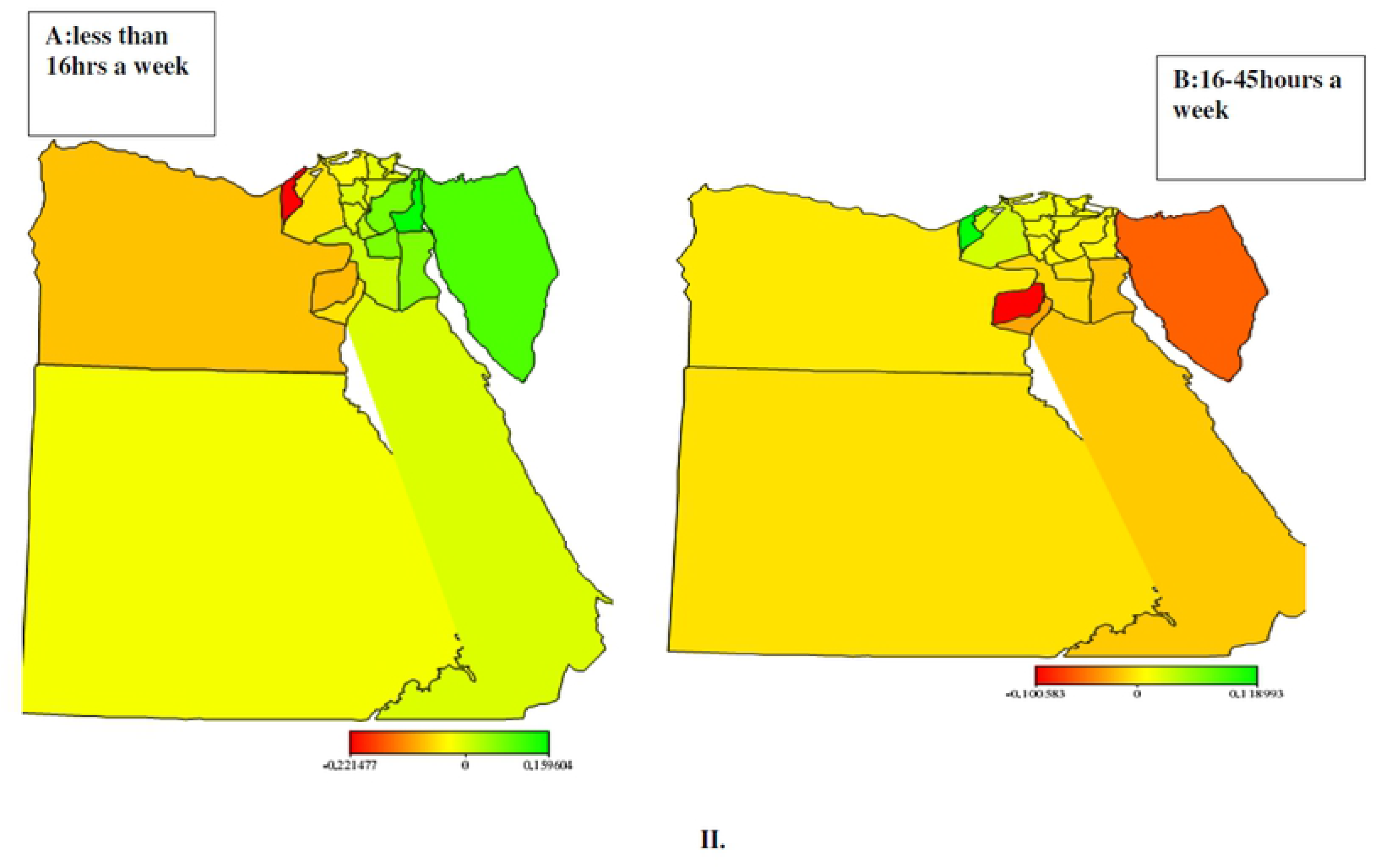
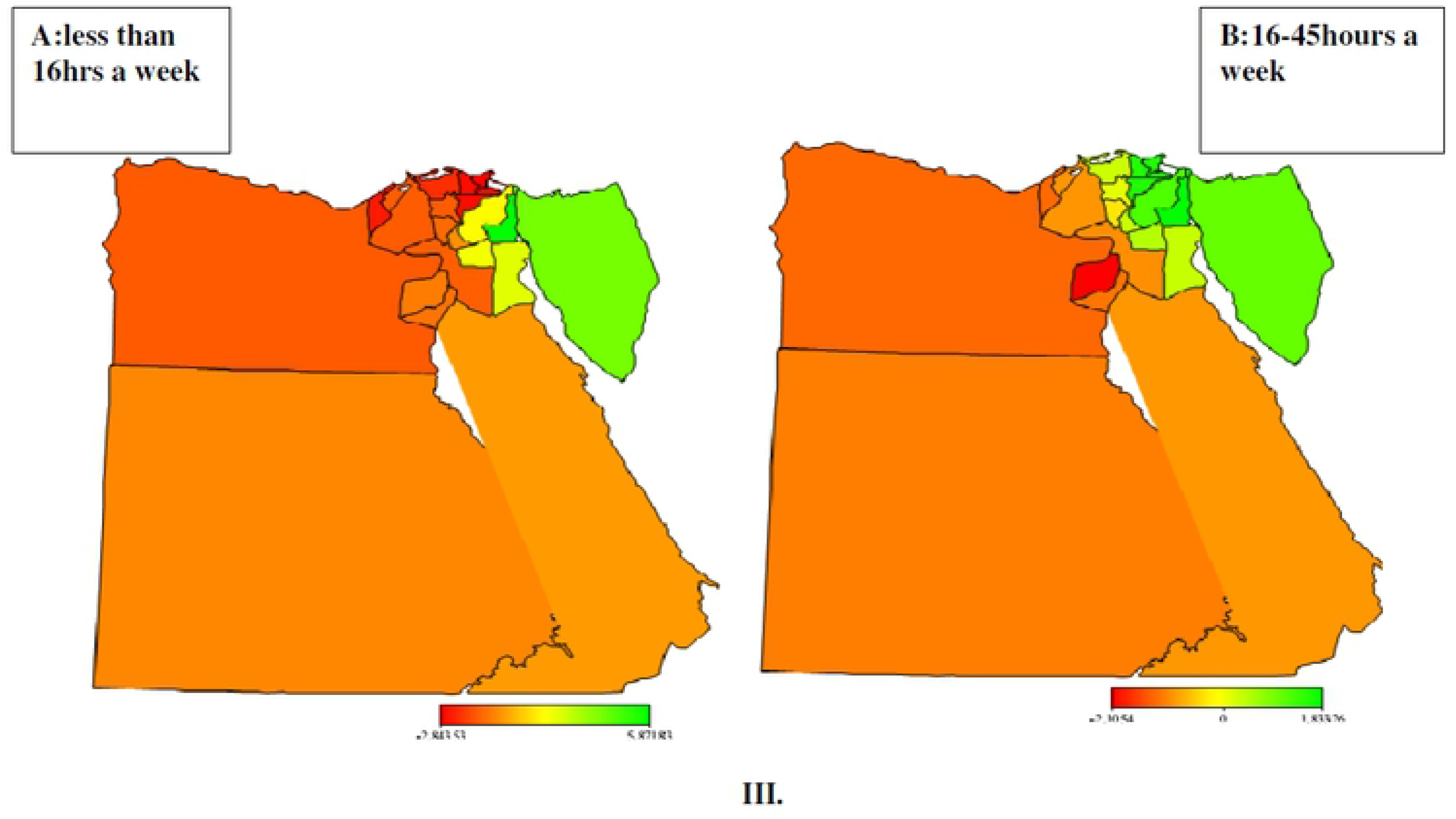
Maps of Egypt showing the spatial effects (posterior OR) on Child labour: I. (A) less than 16hrs a week Vs. (B) 16-45hours a week engagement in Household work, II. (A) less than 16hrs a week vs. (B) 16-45hours a week engagement in wage work, III. (A) less than 16hrs a week Vs. (B)16-45hours a week engagement in Hazard work.

## Discussion

This study emphasises the significance of seeking child protection against engaging in any job or task that could be hazardous, interfere with education, or damage his or her health or physical, mental, spiritual, moral or social development in line with the UN convention. On the note of which we find it imperative to investigate impacts of socio-demographic factors and the associated risk factors on the child labour in Egypt. We reviewed how poverty, declining economic conditions, and rising inflation continue to worsen the situation across various parts in Egypt. To achieve the aim of this study, the UNESCO (2015)’ statistics on the distribution of children aged 5-14 years, of the school age who engages in child labour across different sectors, are examined.

We then cross-classified the economic activities of the children aged 5-17, who are never-married and are of the school age, based on the number of hours worked per week, subject to ILO classifications against the socio-demographic and spatial factors to determine the level of association between these factors and the nature of jobs these children are exposed to. We went further to determine which of the socio-economic and spatial factors and the associated risk factors are likely to expose these children to the risks inherent in the various labours they are engaged.

From the results, we found that gender, age, household size, place of residence, regions, violent discipline method, parental education and survivorships and wealth status of the children are all predisposing factors of the children to the child labour at different working conditions. Specifically, we found that the percentage of male children in non-hazardous jobs; working longer hours are more than female children. However, more than 61% of female children engaging in household jobs spend 16-45hours at work, compared to their male counterparts with about 39% in household jobs. Likewise, female children spend more hours weekly (over 45hrs) in household jobs compared to male children.

Percentage of children living in a medium household and working over 16hrs a week is higher compared to those both in the small and large household. Children living in Upper and Lower part of Egypt, those from the poorest home, those born by women without formal education and those that have experienced both psychological and physical punishments have higher percentage compared to their mates within the respective categories.

Further, we found that children aged 11-15 are more at risk of exposure to child labour than those aged over 15 years. The same goes to the rural children, who have higher chances of exposure than their colleagues from urban cities. Children from the poorer homes, those whose fathers are dead and those subjected to psychological aggression and physical punishment are more likely to be lured into child labour.

Apparently from on our findings so far, one could argue that there is evidence of child labour in Egypt and that socio-demographic and spatial factors greatly predispose majority of the children to it.

## Conclusion

We have presented Bayesian geospatial modelling with multiple imputation models for child labour and violence against child issues in Egypt. These types of epidemiological studies are relatively rare. Most of the previous studies were limited to exploratory analysis of child health only [29], [24] [25].

We have established evidence of the presence of child labour and the impacts of socio-demographic and their associated risk factors on the child labour across Egypt

This study is novel because the association between geospatial factors, socio-demographic factors, and child labour has not been investigated before in Egypt. The findings reveal a significant influence of socio-demographic and economic factors on the child labour and violence against children in Egypt. Significant of these findings is that poverty, neglects, lack of adequate care and exposures of children to various grades of violence are major drivers of child labour across the country. North-eastern region of Egypt has a higher likelihood of child labour than most other regions, while children who live in Delta are more engaged in hazard work. Government is therefore encouraged by the outcomes of this study to work towards protecting the rights of children as enshrined in the United Nations convention and to educate and empower children from the less-privileged families

### Strength and Limitations of this Study

As much as we know, this is the first time study of this kind would be undertaken in Egypt on geospatial factors impacting negatively on child labour. Also, it is the first study to our knowledge which implements such advanced modelling whilst imputing the missing values that usually affect the data of the child labour in developing countries. The study is done in line with the 2014 EDHS, which promotes the assessment of several key aspects of the welfare of Egypt’s children. However, the level of missing observations in the data is a major challenge in this study.

## Author Contributions

Conceived and designed the experiments: KK. Performed the experiments: KK. Analysed the data: KK. Contributed reagents/materials/analysis tools: KK. Wrote the Paper: KK MR.BS. review: KK MR BS MI.

## Supporting Information

S1 Table. Statistics on children work and education in Egypt according to Data from 2013, published by UNESCO Institute for Statistics, 2015

S2 Table. an overview of children’s work by sector and activity in Egypt according to the US Department of labour report, 2016.

S3 Table. Distribution of factors analysed in child labour in Egypt (DHS 2014)

S4 Associate Risk Factors with Child Welfare including Child Labour and Violence against Child issues in Egypt (DHS 2014).

S5 Missing values among the risk factors

S1 Data EDHS data 2014 on child labour

## Footnotes

1 The Kish grid gives a procedure of selection. The expression “Kish grid” comes from the name of Leslie Kish, the Hungarian born American statistician. Kish was one of the world’s leading experts on survey sampling.

